# inPhase — A simple, accurate and fast approach to determine phase diagrams of protein condensates

**DOI:** 10.1101/2024.10.02.616352

**Authors:** Anatol W. Fritsch, Juan M. Iglesias-Artola, Anthony A. Hyman

**Affiliations:** Max Planck Institute of Molecular Cell Biology and Genetics, 01307 Dresden, Germany; Center for Systems Biology Dresden, 01307 Dresden, Germany; Cluster of Excellence Physics of Life, Technische Universität Dresden, 01307 Dresden, Germany

## Abstract

Protein phase separation has become a widely studied phenomena in biology with implications in cell metabolism and disease. The study of phase separating proteins often relies on the precise determination of their phase diagrams. These phase diagrams give information on the protein concentration required for condensate formation and the respective concentration inside the condensate at defined external conditions (temperature, salt, pH). However, it has so far often proven difficult to accurately measure phase diagrams. Here, we report a method that is based on mass and volume conservation and defined reaction volumes, which we call inPhase. We can use this method to determine accurate values for both dilute and condensed branch protein concentrations. With this information we can produce accurate phase diagrams. We compare our method to the widely used quantitative fluorescence approach and find that it underestimates the partition factor into condensates at least two-fold for FUS and PGL-3. The accessibility of our method opens the possibility for the thermodynamic assessment of entire protein families, generating sufficient quantitative data for testing theory, and for the screening of drugs in pharmaceutical research.

## Introduction

Cells carry diverse metabolic functions that require distinct biochemical and biophysical environments. Recent advances in cell biology show that non-membrane bound compartments can generate distinct environments and that they are relevant for the functioning of a cell. These compartments, referred to as biomolecular condensates, can in some cases form through the process of phase separation, where the interactions among certain proteins (with or without nucleic acids) are thermodynamically more favourable than with the surrounding cytosol^1–3^. Protein condensates are present in a wide variety of cells where they perform fundamental functions in development (e. g. P granules in *C. elegans*^4,5^); ageing and stress (e.g. stress granules^6,7^); and in diverse cellular processes (e.g. transcription^8,9^).

Perhaps the most pressing question is the following: How have proteins evolved to carry out precise and reproducible phase separation inside cells? We understand well how synthetic polymers phase separate. But these are simple compared to proteins, the amino acid composition of which results in incredible complexity compared to synthetic polymers.

The propensity of a protein to phase separate is ultimately governed by its sequence and its environment^10,11^. Environmental factors have implications, for example, in the embryogenesis of *C. elegans* or on self-preservation under low pH or high temperatures in yeast. In general, temperature, salt concentration and pH are biologically significant external control parameters of thermodynamic phase separation^12–16^. From these considerations, it follows that to understand the phase separation behaviour of a given protein and its potential function and physiological relevance, its phase diagrams must be constructed with respect to the relevant variables under study. Only when we understand the relationship between amino acids and phase separation properties can we hope to modulate phase separation. Without such accurate data it is hard not only for biochemists, but also physical chemists and physicists to develop quantitative models.

A phase diagram depicts a binodal curve that contains information on the protein concentration in the dilute (protein poor) and condensed (protein rich) phase (see Fig. 1a) under the given conditions. As is common in nascent fields, accurate and widely accessible quantitative methods to determine phase diagrams are still lacking. While dilute phase protein concentrations, or saturation concentrations, have been reported in numerous studies^10,12^; condensed phase concentrations have been more complicated to measure^17–19^, and to our knowledge methods do not exists to determine these simultaneously.

**Figure 1.**
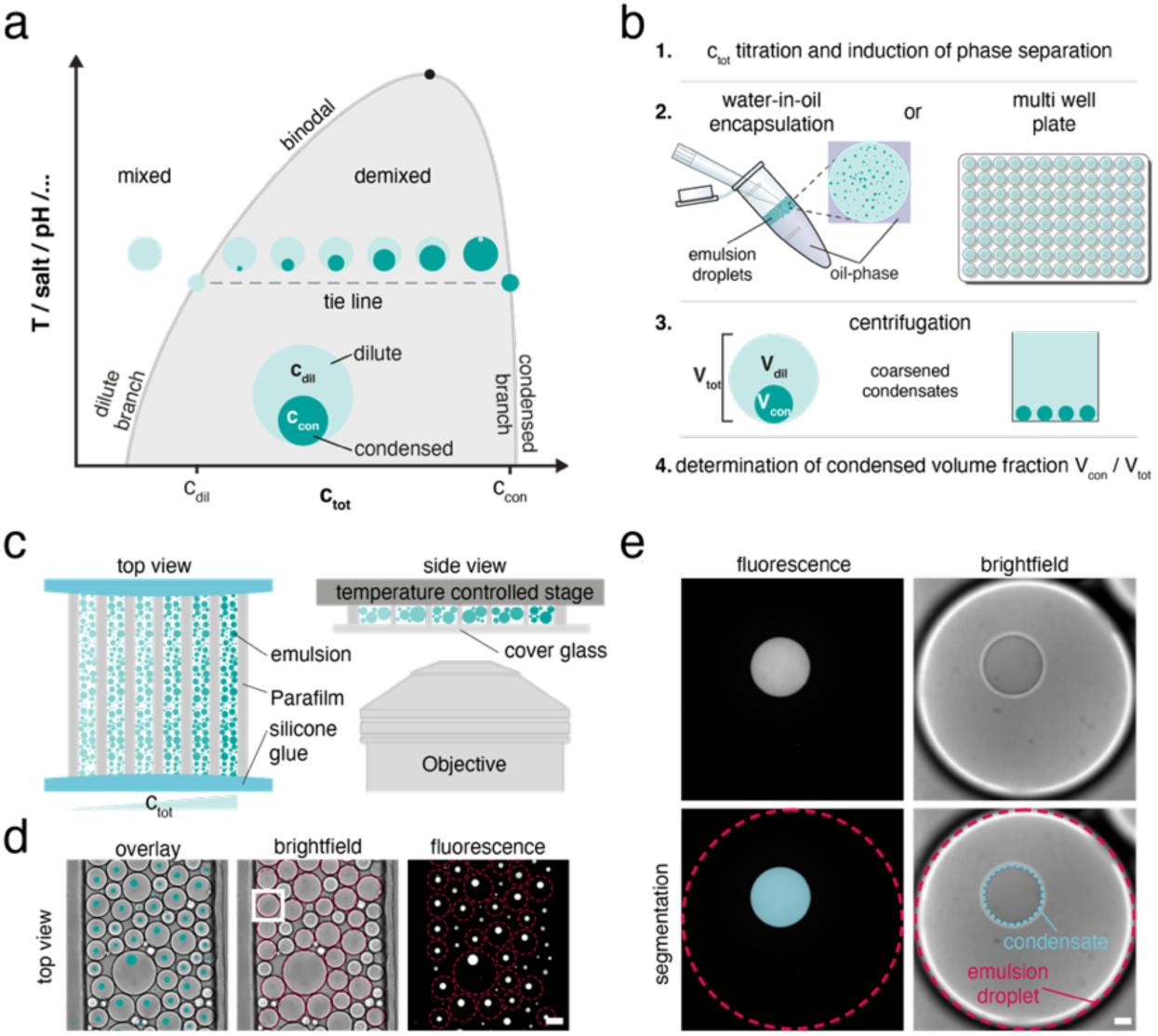
Workflow of inPhase. **a**, Schematic phase diagram showing the binodal, consisting of a dilute branch and a condensed branch. A pair of c_dil_ and c_con_ concentrations is connected via a tie line. **b**, Experimental protocol consisting of four consecutive steps. **c**, Schematic of a titration series in water-in-oil emulsions mounted on a temperature-controlled stage. **d**, Images of emulsions droplets containing PGL-3 condensates for one c_tot_ (scale bar 100 µm). Left panel is an overlay of the brightfield (middle) and fluorescence (right) images. Dashed red circles show the segmentation of the emulsion droplets. **e**, Magnified view of an emulsion (white square in panel d) and its condensate. Top panels show the fluorescence and brightfield images. The bottom panels show the fluorescence (left) and brightfield (right) based segmentation of the condensate and emulsion. Scale bar: 10 µm.

In this study we develop a fast and accurate approach, called inPhase, to produce phase diagrams and thereby to determine both dilute and condensed phase concentrations of binary protein mixtures. Our method is based on the fact that the condensed phase volume changes linearly with protein concentration^20^. This principle can be used to estimate the protein concentration in the dilute and condensed phase. The method does not rely on fluorescent tags, is independent of a specific setup and only relies on the knowledge of the condensed phase volume in a container of defined volume.

To showcase our method, we measured salt and temperature dependent phase diagrams of two well studied proteins, fused in sarcoma (FUS) and *Caenorhabditis elegans’* P granule protein PGL-3. We demonstrate how rapid construction of these phase diagrams gives novel insight into the phase behaviour of these proteins. The phase diagrams allow us to select conditions for temperature shift experiments to test the reversibility of phase separation. Our results show that quantitative fluorescence intensity-based approaches underestimate the partition factor of the proteins into the condensed phase by at least two-fold.

## Results

### inPhase concept

We assume that when a protein phase separates in a closed system, the total volume of the system, *V*_*tot*_, is conserved and it can be expressed as the sum of the volumes of the dilute phase and the condensed phase (*V*_*tot*_ = *V*_*dil*_ + *V*_*con*_, Fig. 1b). In this closed system, the total protein mass is conserved (*m*_*tot*_ = *m*_*dil*_ + *m*_*con*_) and given that the mass equals the volume times the concentration (*m* = *V* · *c*), we can express the previous equation as *V*_*tot*_ *c*_*tot*_ = *V*_*dil*_ *c*_*dil*_ + *V*_*con*_ *c*_*con*_. Rearranging the volume and mass conservation equations leads to:

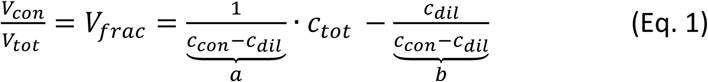

Thus, we see that the ratio *V*_*con*_⁄*V*_*tot*_, referred to as the volume fraction *V*_*frac*_, changes linearly with the total protein concentration, *c*_*tot*_, of the system (Extended Data Fig. 1). A linear regression to a total concentration series allows us to determine both *c*_*dil*_ = −*b*/*a* and *c*_*con*_ = (1 − b)/a. It therefore follows that we can determine *c*_*con*_ and *c*_*dil*_ at defined external conditions (temperature, salt, pH) by preparing samples at different total protein concentrations, *c*_*tot*_, and by measuring the volume of the condensed phase, *V*_*con*_, in a closed system of defined volume, *V*_*tot*_. In principle, this allows us to measure phase diagrams for any ordinary binary system.

To experimentally set a closed environment of defined volume, we used a water-in-oil emulsion system or 384 well-plates (Fig. 1b). An experiment consists of a dilution series of the protein of interest at phase separating conditions (Fig. 1c). For each dilution we determine the volume fraction *V*_*frac*_ of the condensed phase. We fit Eq. 1 to this data and obtain both *c*_*con*_ and *c*_*dil*_. In the emulsion system *V*_*frac*_ can be measured by image segmentation of condensates and emulsion droplets (Fig. 1d, e). In the well plates we only need to segment the condensed phase because the total volume is known. The presence of tagged-protein is not necessary for the segmentation of the condensates but facilitates the process. Repeating this procedure at different conditions (salt, pH, temperature) results in a binodal curve (Fig. 1a). We calibrated our image analysis pipeline with fluorescent polystyrene beads of known size and conclude that we can accurately determine the volume of the condensed phase (Extended Data Fig. 2).

### inPhase returns accurate c_con_ and c_dil_ values

We used PGL-3, a protein component of P granules found in the germline of *C. elegans,* as an example to apply our method. We observed that the linear regressions are in good agreement with our experimental data (Fig 2a, Extended Data Fig. 3a-g). As expected, the condensate phase volume increases linearly with increasing protein concentration as visually shown in Fig 2a (lower panel).

**Figure 2:**
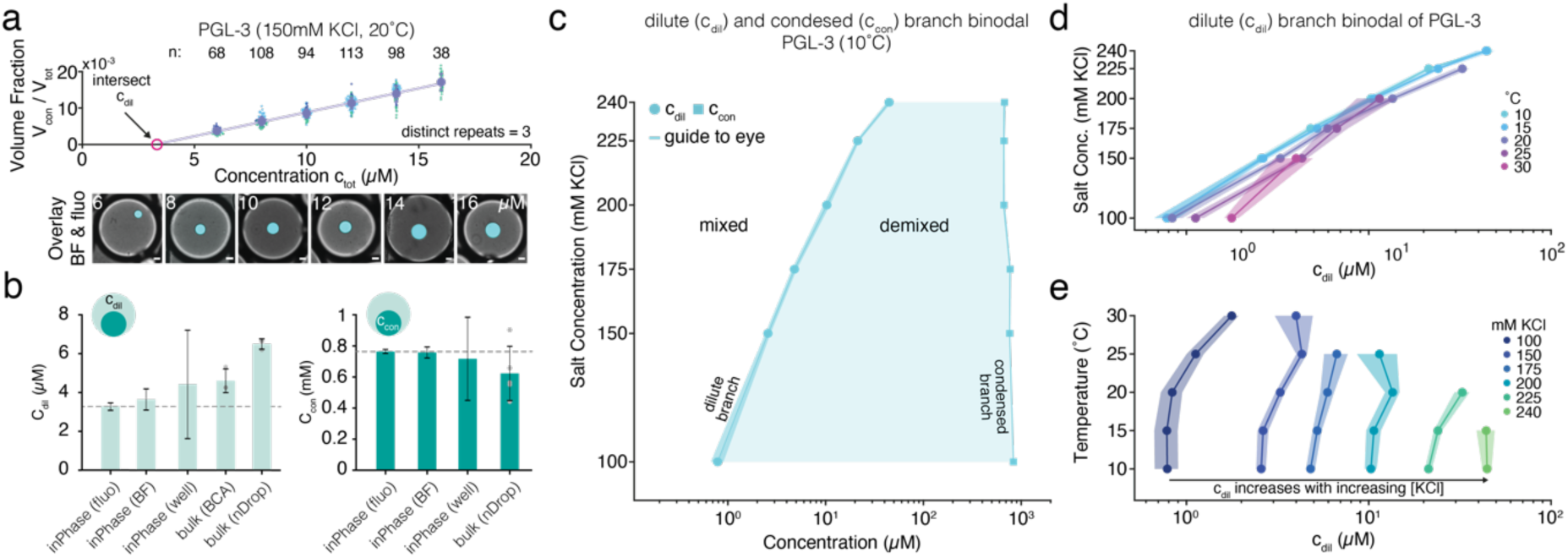
Salt and temperature dependence of c_dil_ and c_con_ for phase separated PGL-3 protein. **a**, Experimental data showing the linear relationship between condensed volume fraction V_frac_ and total protein concentration c_tot_ for PGL-3 (5% PGL-3::GFP). The scattered data points indicate the V_frac_ of individual emulsion droplets (color-coded by distinct repeats), big data points show the mean of the V_frac_ of individual emulsion droplets at a defined concentration. The error bars represent the SD of the individual emulsion droplets. Eq.1 was fitted to all the individual emulsion droplets V_frac_ using a robust b-square fitting routine. The lower panel shows images of emulsions with increasing size of condensates for increasing c_tot_. **b**, Comparison of c_dil_ and c_con_ derived via inPhase using fluorescence (fluo) or brightfield (BF) for segmentation (in emulsions or multiwell plate) versus bulk measurements via Pierce BCA (BCA) and absorption measurements using Nanodrop (nDrop). The dashed line indicates inPhase (fluo) value as reference. Independent measurements are shown in gray. For inPhase methods, the error bars show the SD derived by error propagation of the SD of the fit parameters in Eq. 1. For the BCA and absorption measurements the error bars show the SD of three independent measurements. **c**, Salt dependent phase diagram of PGL-3 at 10°C showing the dilute (c_dil_, closed circles) and condensed (c_con_, closed squares) branches. The shaded region along the line shows the estimated SD. **d**, **e**, Dilute branch concentrations for salt (d) and temperature (e) changes of PGL-3. The shaded areas represent the SD obtained by error propagation from the SD of the fit. For the exact sample size at each data point see Supplementary Table 1.

We then compared the measured dilute and condensed protein concentrations using the different modes of inPhase (fluorescence, brightfield and well-plate) with measurements obtained using light absorbance (nanoDrop) and the BCA (bicinchoninic acid) assay (Fig. 2b). Our results show that our method returns accurate values for both the dilute and the condensed phase protein concentrations when compared to light absorbance and BCA measurements. While the well-plate assay in the current iteration is less precise, the average value of the other measurements is within the error bars. The discrepancy of the absorbance measurement for the dilute phase to all other measurements can be explained by the low protein concentration, which lies at the detection limit of the used light absorbance device.

### Acquiring phase diagrams with inPhase

Performing inPhase at a defined temperature and at different salt concentrations results in a salt dependent binodal (Fig. 2c). As expected, we find that *c*_*dil*_ increase for higher salt concentrations, while *c*_*con*_ is not considerably affected. We then measured salt dependant binodals at different temperatures (Extended Data Fig. 3h, i). We observe that temperature has a smaller effect on *c*_*dil*_ than does KCl concentration in the measured range (Fig. 2d). This effect can be observed more clearly when plotting the temperature dependent dilute branches at different salt concentrations (Fig. 2e). In this diagram the change of *c*_*dil*_ with temperature is small (as shown by rather vertical dilute branches) while the change in *c*_*dil*_ with KCl concentration is clearly visible as a shift in the location of the dilute branch along the protein concentration axis.

We then proceeded to measure the different temperature and salt dependent phase diagrams of the widely used fused in sarcoma (FUS) protein (Extended Data Fig. 4). In contrast to PGL- 3, the phase diagrams of FUS show a strong dependence of *c*_*dil*_ to temperature and a moderate dependence to salt concentration. We then harboured this knowledge to further assess the validity of our *c*_*dil*_ estimates and the reversibility of the condensation process.

### Testing binodals with dynamic temperature shifts

An easily accessible experimental value to address reversibility is temperature. We therefore chose FUS for these experiments owing to its strong dependence of *c*_*dil*_ on temperature.

From the temperature phase diagrams of FUS at different salt concentrations, we can determine a protein concentration that allows us to move in and out of the demixed region within a reasonable temperature range (Fig. 3a). We prepared samples at 100, 150, 200 or 300 mM KCl that contained 2.4, 4.8, 7.9 or 15.9 µM of FUS, respectively. From the phase diagrams we expect that all the samples will phase separate at similar temperatures, termed condensation temperature *T*_*cond*_. Starting from 30 °C we reduced the temperature to 10 °C as depicted in Fig 3b (upper panel). As expected, all protein mixes started to phase separate around the predicted temperatures between 17 and 23 °C (indicated by the black region in graph in Fig 3b). The slightly lower *T*_*cond*_ for 300 mM KCl compared to 100 mM KCl is also reflected in a later entry into the phase separated region in the dynamic experiment. While the higher salt concentrations dissolve all condensates after heating to the initial temperature, small remnants are still visible after reheating for the lower salt concentrations (orange dashed lines in Fig 3b lower). Furthermore, the volume fractions *V*_*frac*_ seem to reach a plateau quickly after entry into the two-phase region and are in line with our reported values expected for these external parameters.

**Figure 3:**
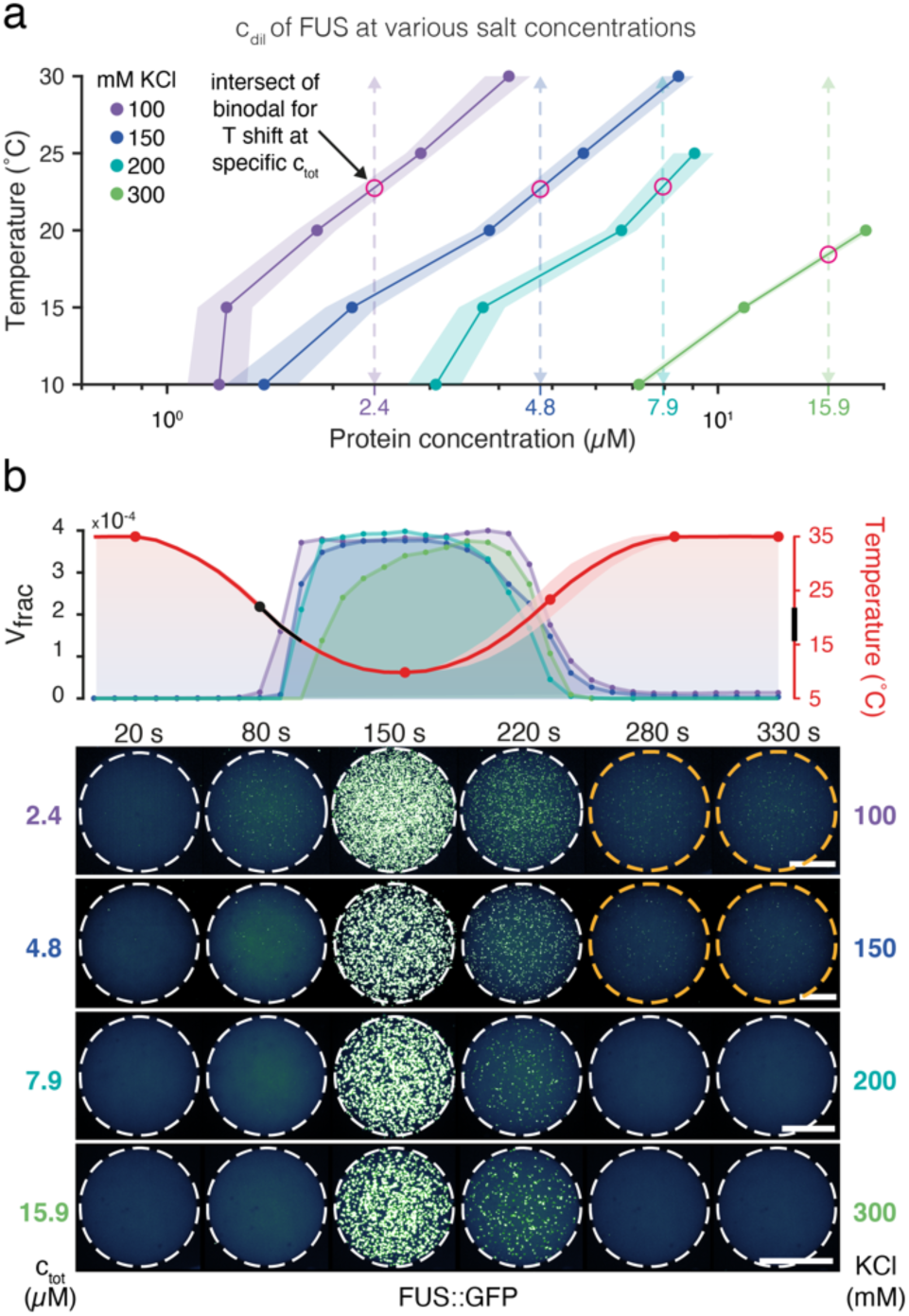
Tuning the reversible phase entry of FUS by fixing salt and protein concentrations. **a**, Dilute branch of the binodal (c_dil_) for FUS versus temperature at various salt concentrations. Red circles demark the condensation temperature for each condition (dashed arrows indicate the temperature shift performed in **b**). The shaded regions show the SD obtained by error propagation of the SD of the fit parameters. For sample size see Supplementary Table 2. **b**, Timeseries of temperature quenches from 35°C to 10°C and back to 35°C. Upper panel: Experimental data of the volume fraction and temperature for four different salt and protein concentrations of FUS (10 s resolution). Black line indicates temperatures with first recorded condensates. Lower panel: Maximum intensity projections of 3D image stacks at specific timepoints (indicated by the red dots on the temperature curve, also Supplementary Video 1-4). White dashed circles indicate emulsions and orange circles indicate conditions where condensates or aggregates remained after reheating. The shaded region of the temperature curve shows the SD of the temperature measurements of the different experiments. Scale bar: 50 µm.

We observe that our measured dilute-branches are in good agreement with the dynamic temperature shifts presented. This showcases the benefit of such a detailed understanding of a protein system for further experimental work on the dynamics of phase separation.

### Calculation of partition factors

The partition factor (PF) can be defined by the ratio of the condensed- over dilute-phase concentration (*PF* = *c*_*con*_ / *c*_*dil*_). It is often used in condensate research to characterise the increase in concentration of protein in the condensed phase. It is common to determine the partition factor by comparing fluorescence intensity inside the condensate against the intensity in the dilute phase. In contrast, our method allows us to directly determine the partition factor by using the presented phase diagrams.

For FUS we observe partition factors in the order of 10^2^-10^3^ that depend on the temperature and salt concentration (Fig. 4a left panel). The partitioning increases with decreasing salt concentrations until 100 mM KCl after which this trend starts to reverse. The partition factor of FUS exhibits a minimum that shifts towards lower temperatures for increasing salt concentrations. This minimum corresponds to the onset of an increased concentration in the condensed phase (Extended Data Fig. 4). We speculate that this increased concentration is a consequence of a rearrangement of the protein with temperature inside the phase. For the partition factor of PGL-3 we observe a small influence of temperature while salt concentration can change it by an order of magnitude, between 10^1^-10^2^ (Fig. 4a right panel). This trend is only broken for higher temperatures and salt concentrations where a sharp increase in the partition factor takes place as is the case for FUS condensates.

**Figure 4:**
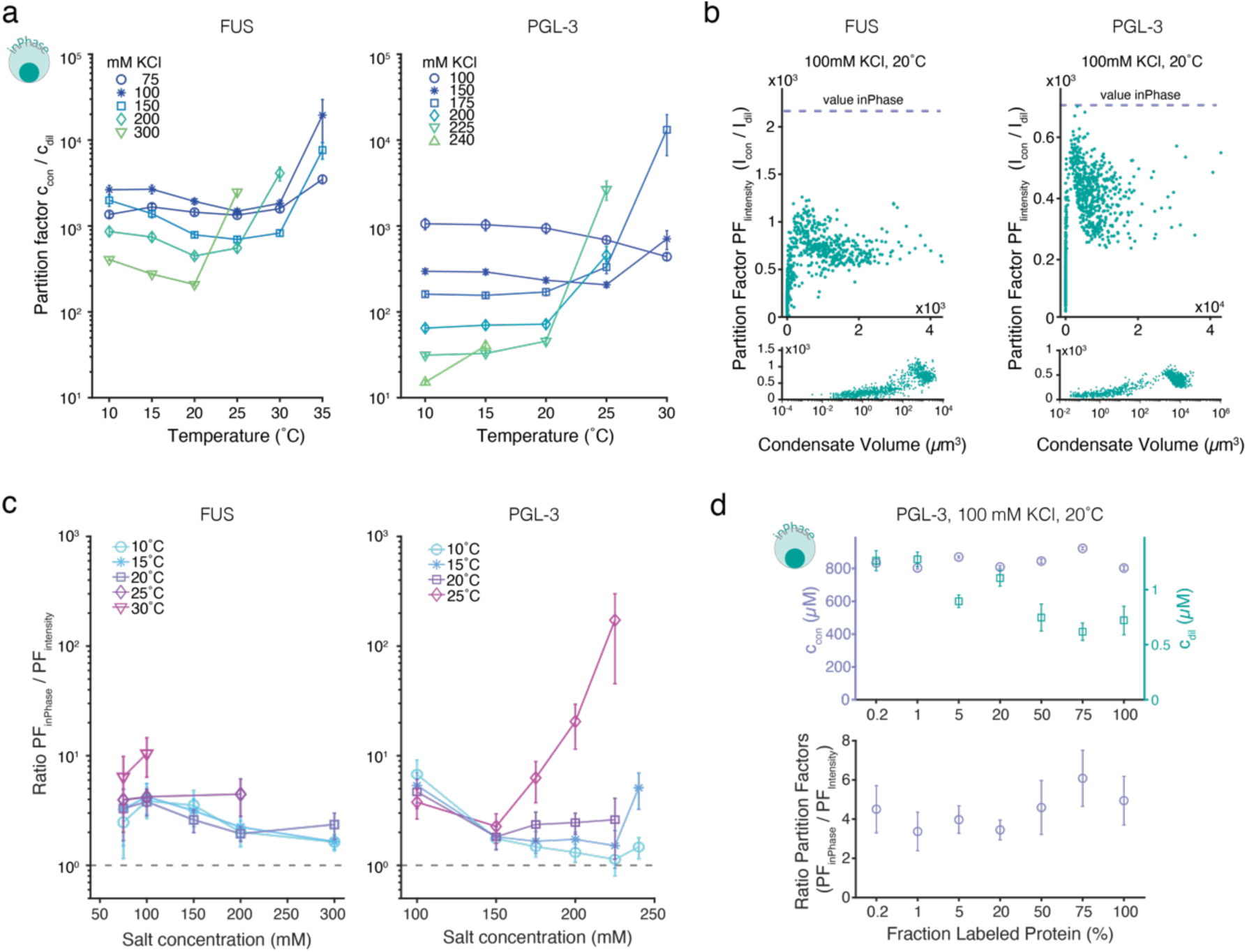
Effect of temperature and salt on the partition factor and comparison to quantitative fluorescence for FUS and PGL-3. **a,** Partition factor PF_inPhase_ (c_con_ / c_dil_) derived via inPhase method for various temperatures and salt concentrations for FUS::GFP (left panel) and PGL-3 + 5% PGL-3::GFP (right panel). Error bars show the SD obtained by error propagation of the SD of the fit parameters **b**, PF_Intensity_ of individual condensates as a function of their volume for FUS::GFP (left panel) and PGL- 3 + 5% PGL-3::GFP (right panel). Upper and lower panels contain the same data with the lower panels using a logarithmic x-axis (dashed lines indicate PF_inPhase_ for the same data-set). **c**, Ratio PF_inPhase_ /PF_Intensity_ for various salt concentrations and temperatures for FUS (left panel) and PGL-3 (right panel). Error bars show the SD obtained by error propagation of the SD of the fit parameters **d,** Partition factor (PF) as a function of the amount of labelled PGL-3 protein fraction. The upper panel shows the c_con_ (purple) and c_dil_ (green) calculated using inPhase. The lower panel shows the ratio of PF_inPhase_/PF_Intensity_ at different GFP percentages. PFIntensity underestimates the partition factor. Error bars show the SD obtained by error propagation of the SD of the fit parameters. For the sample size see Supplementary Table 1 and 2.

### Comparison of partition factors to fluorescence-based quantification

Next, we compared our calculated partition values to the ones obtained using quantitative fluorescence intensity. Given that in this instance we used fluorescence to segment the condensate volume, we can use the same data to directly compare both methods (*PF*_*intensity*_ via fluorescence intensity, *PF*_*inPhase*_ via inPhase). As expected, we report a strong dependence of the intensity-derived partition factor *PF*_*intensity*_ on condensate size for both FUS and PGL-3 (Fig. 4b). Above a condensate volume of approximately 100 µm^3^ the partition factor *PF*_*intensity*_ levels off to a plateau-like value that is still lower than the value we calculated using inPhase (purple dashed line). We then further estimated the *PF*_*intensity*_ values by using condensates that lie within the plateau region.

In general, the partition factor using fluorescence intensity underestimates the protein partitioning compared to our method (Fig. 4c). The difference between *PF*_*intensity*_ and *PF*_*inPhase*_ is more prominent for FUS than for PGL-3, and at least 2-fold different over the entire dataset. This effect is likely caused by an erroneous estimation of the fluorescence intensity inside the condensate phase that is caused by limitations of the optical system as well as by a different chemical environment inside the phase.

To address the possible effect of fluorophore quenching on the calculation of the partition factor, we compared *PF*_*intensity*_ to *PF*_*inPhase*_ at different fractions of labelled PGL-3 (Fig. 4d). Our measurements show that quantitative fluorescence microscopy underestimates the partition factor independently of the label fraction used. Decreasing the label fraction of PGL- 3 did not result in a considerable reduction of the difference between the methods. This indicates that fluorophore quenching by the fluorophore up-concentration in the condensed phase is not the main cause for the difference between the methods.

We then used the calculated *c*_*dil*_ and *c*_*con*_ values to understand how the labelling percentage affects the phase separation process. Our results show that *c*_*con*_ remains relatively constant when preparing condensates with 0.2% to 100% of labelled PGL-3. On the other hand, half of the protein concentration *c*_*dil*_ is required to achieve phase separation when using 100% labelled protein compared to 0.2% of PGL-3. This indicates that GFP increases the propensity of PGL-3 to phase separate under the conditions studied.

Overall, we show that inPhase can accurately produce protein phase diagrams that contain the protein concentration in the dilute and condensed phase using fluorescence or brightfield microscopy. The temperature shift assays at different salt and protein concentrations show that we can determine the condensation (cloud point) temperature from the phase diagrams. We have additionally shown that our method is more robust in measuring the partition factor when compared to quantitative fluorescence microscopy. Finally, we have tested if fluorescent protein tags affect condensate formation and conclude that PGL-3 condensates formed with 100% labelled protein require a lower protein concentration for phase separation.

## Discussion

Currently, only a few protein phase diagrams have been reported that present both the dilute and condensed phase protein concentrations^16,17,19,21–24^. In general, phase diagrams contain rich information that can be used to assess protein interactions, the molecular grammar of phase separation and to validate current theoretical approaches^24–26^. Despite their central role, current methods to acquire phase diagrams are still not widely accessible. inPhase offers a powerful method to address this challenge. We showed that we can generate accurate phase diagrams at defined temperature and salt concentrations for PGL-3 and FUS with a minimal amount of protein. Based on these phase diagrams we can perform dynamic temperature experiments to assess the reversibility of phase separation and to test the accuracy of the measured c_dil_ values.

We have shown that inPhase does not necessarily rely on fluorescent protein tags and that it can be performed in well-plates for future screening assays e.g., to quantify the effect of small molecules in drug research. Our method is also transferable to microfluidic-based emulsion systems, that allow to change protein concentrations during an experimental run. Our method can also be extended to cellular systems as for example to one-cell embryos of *C. elegans* (see Extended Data Fig. 5). This because a population of cells or embryos inherently exhibits a heterogeneous protein concentration and their total volume as well as the condensed phase volume of interest can be measured. Furthermore, ternary mixtures with additional RNA could also be addressed (see Extended Data Fig. 6). For multicomponent mixtures the corresponding c_con_ is however an apparent measure that needs further investigation.

inPhase is based on the mass-conservation law and the assumption of volume conservation to measure concentrations of the dilute and condensed phase of phase separating proteins. The underlying principles and techniques make this method accessible to the research community as it relies on commonly available equipment (fluorescent or bright field microscope), modest computational power and low amounts of protein. Our approach does not necessitate the use of labelled protein, which not only reduces the time required for cloning and protein purification but could also remove artefacts associated with the use of fluorescent tags (see Fig. 4c). Furthermore, the accuracy of inPhase has the potential to increase reproducibility in the protein phase separation community and it offers the possibility to test theoretical models.

Other methods to study the dilute or condensed branch concentrations include more elaborate bulk measurements^27–29^, measurements of optical density^18,30^, microfluidic systems^31,32^, capillary flow experiments^33^, or fluorescence correlation spectroscopy (FCS)^22^. Bulk approaches are accurate but require large amounts of protein that are often not available. Optical density measurements provide a reliable way to determine the condensed branch concentrations as well as a limited possibility to be used in an *in vivo* setting; however, they do not provide dilute branch concentrations, need dedicated equipment, and are potentially difficult to be used for screening. Finally, FCS can also be used to measure phase diagrams but imposes constraints on sample preparation and inherently uses fluorescence as a readout.

An accurate condensed phase segmentation is crucial for our method. In its current form, our segmentation assumes that condensates are spherical. However, condensates can flatten due to gravity and wetting. We have reduced wetting by using water-in-oil emulsions and low-binding well-plates. Nonetheless, condensates above a certain size tend to flatten. This will result in an overestimation of condensed phase volume. The effect becomes apparent for PGL- 3 samples prepared at high concentrations. However, this artefact can be overcome by performing 3D segmentation of the condensates.

A typical measurement of a temperature dependant phase diagram requires approximately one hour to be completed. We therefore corroborated that in this timeframe the properties of condensates do not change considerably (Extended Data Fig. 7). In addition, this shows that this method can be used to study the time dependence of condensate properties. For PGL-3 we show an increase of c_dil_ and a decrease in c_con_ over time, while FUS values are rather constant in the timeframe reported.

Overall, we propose inPhase as an efficient new method for measuring phase diagrams, which we believe will contribute to the study of phase separating proteins. We consider that the concept at the core of this method is readily accessible to a wide community and that the experimental workflow, performing a protein titration, can be readily adapted. Our method can be used to study the effect of protein modifications, to further elucidate the molecular grammar of phase separation, and to validate theoretical models.

## Data availability

The summary data (mean ± SD) of the phase diagrams are available in Supplementary Table 1 and 2. All movie and image raw data (including the processing steps) are archived on magnetic tape at the Max Planck Institute of Molecular Cell Biology and Genetics, Dresden, and can be made available upon reasonable request.

## Code availability

The MATLAB code used to analyse the microscopy images can be accessed at the GitLab space of the Max Planck Institute of Molecular Cell Biology and Genetics, Dresden: https://git.mpi-cbg.de/fritsch/inphase-code-repository

## Methods

### Preparation of emulsions

Protein solutions were prepared from stocks that are frozen at −80°C. The protein stock solutions were cleared from potential aggregates using small volume centrifugal filters (UFC30VV25, Merck, Germany). The protein concentrations were determined via extinction coefficient measurements at 280 nm (NanoDrop, Thermo Fisher Scientific, USA). Next, the desired fraction of labelled protein was prepared (main experiments 5% GFP labelled protein for PGL-3, 100% for FUS). The protein concentrations were adjusted using the same salt concentration as the protein stock (300mM KCl and 25mM HEPES for PGL-3 and 500 mM KCl and 50mM HEPES for FUS; both 1 mM DTT at 7.4pH). In the final step the salt concentration was reduced to the desired value, using a buffer containing no salt (HEPES and 1 mM DTT at 7.4 pH) to trigger phas separation. The protein solution was then immediately encapsulated into a water-in-oil solution. Encapsulation was performed using twice the amount of PicoSurf oil (2% (w/w) in Novec 7500, Sphere Fluidics, UK) compared to the protein solution. Here, we add the oil to the protein solution in standard 100µl PCR tubes and the solution is agitated using a 20µl pipette and 200 µl tips until the desired size distribution of water-in-oil solutions is reached. To coarsen the phase separated condensates within the emulsion drops we used a centrifugation step (200-400 g for 4 min) at the desired temperature of the first data point. This will lead to a single condensate per emulsion droplet in most emulsion droplets. A titration series of the total protein concentration was then mounted on a temperature-controlled microscope stage.

### Sample loading and imaging procedure for emulsions

For mounting we use a custom-made temperature-controlled microscope stage based on Peltier elements connected to a sapphire microscope slide (similar to Mittasch et al.^34^). This allows for fast and precise temperature changes both below and above room temperature. To allow for parallel imaging of titration series we generate lanes of Paraflim (Bernis, USA) sandwiched between the sapphire slide and a standard cover slide (22×22 mm, 170µm, Menzel, Germany; also see Fig. 1c). Emulsion solutions are loaded into the individual lanes by pipetting the emulsion. Each lane is sealed using addition curing silicone (Picodent twinsil speed, Picodent, Germany) to minimize evaporation and movement of the water-in-oil emulsion. Sample imaging was performed via CellSens software (Olympus, Japan) on an Olympus IX83 microscope connected to a Yokogawa W1 SoRa spinning-disc system (Yokogawa, Japan) and a Hamamatsu Orca Flash v4 sCMOS camera (Hamamatsu, Japan) using a 40x air objective (UPLXAPO, 0.95NA, Olympus, Japan). For each sample and each parafilm lane, a multi-tile brightfield image and a corresponding multi-tile 3-dimensional fluorescence stack was recorded using a piezo stage (Mad city labs, USA). The main temperature experiments started at 10°C and increased the temperature by 5°C steps. Between consecutive temperatures we added 5 min of equilibration time before imaging.

### Image analysis

Images were processed by a custom-written MATLAB (The Mathworks, USA) script using OME Bio-Formats^35^ for image and metadata handling. The brightfield images of each lane were used to detect the contours of the water-in-oil emulsion droplets. The location as well as the volume of each emulsion droplet was determined assuming that the height of the Parafilm chambers is of approximately 100 µm. We assumed an ellipsoid shape for emulsions with a radius bigger than 50 µm. Corresponding positions of each emulsion droplet are further analysed in the fluorescence image stack. For each emulsion droplet the volume of the condensed phase is determined via image registration as well as further parameters like the intensity of the dilute and condensed phase. In brief, after image projection and basic image filtering, we use an intensity gradient approach to find a good estimate for the outline of the condensates. We then assume sphericity of each condensate to calculate its volume. For large condensates the spherical assumption might introduce overestimation of the actual volume since they can flatten out due to gravity. To get valid estimates for the dilute phase intensities we first dilate the masks of the condensed phase and remove them from the mask of the emulsion droplet. This helps to minimize the influence of intensity values close to the margins of the condensate phase. Furthermore, we can also use the brightfield images to estimate the volume of the condensed phase. The image analysis pipeline was calibrated using monodisperse fluorescent beads of known size immersed in glycerol solution to mimic the refractive index difference found for condensates in buffer (Extended Data Fig. 2).

### Calculation of dilute and condensed branch concentrations

To get to the form of Equation 1 we assume volume and mass conservation in the reaction container. While there could be processes that lead to a change of total volume in case of a phase transition in our case of protein phase separation the actual volumes of the condensed phase are orders of magnitude smaller than the total reaction volume and thus even in such a case the linear form will be valid for small volume fractions *V*_*frac*_ = *V*_*con*_⁄*V*_*tot*_).

The conservation reads as *V*_*tot*_ = *V*_*dil*_ · + *V*_*con*_ for the volume and as *V*_*tot*_ · *c*_*tot*_ = *V*_*dil*_ · *c*_*dil*_ + *V*_*con*_ · *c*_*con*_ for the mass given that the mass equals the volume times the concentration (*m* = *V* · *c*). Converting the mass conservation to 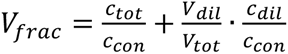 and substituting the volume conservation as 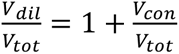 leads to 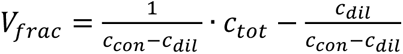 (Equation 1). The volume fraction data for each total concentration was pooled from all repeats and used for linear regression of the concentration series in *c*_*tot*_. Using the linear form *V*_*frac*_ = *a* · *c*_*tot*_ + *b* we get 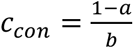 and 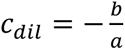 according to Equation 1. Via the confidence interval of the fit and error propagation we can determine an error for the dilute- and condensed-phase concentrations.

### Multi-well plate assay

We load equal amounts of 20 µl sample volume for the total protein concentration titration in 384 well plates (PhenoPlate, PerkinElmer, USA). The well plates were chosen due to the superior non adhesive properties that allow to assume spherical shape of the condensates. Well plates are centrifuged at 200 g for 10 min with low acceleration and deceleration of the rotor to minimize coarsening into one condensate. The segmentation routine used for the emulsion-based approach was calibrated for the well plates and used to determine the volume fraction of the condensed phase for each total concentration. We then use equation 1 to derive *c*_*dil*_ and *c*_*con*_ and the respective errors.

### Bulk concentration measurements

Centrifugation of a phase separated suspension provides us with the supernatant containing the dilute phase and a condensed phase at the bottom of a PCR tube that can both be used for the bulk measurements when using large amounts of sample. We could use our measured volume fractions of condensed phase for given external parameters to arrive at volumes of condensed phase suitable for pipetting. We used the Pierce BCA protein assay kit (ThermoFischer, USA) together with a TECAN Spark 20M plate reader (TECAN, Swiss) as reference to determine the dilute branch concentration. For each measurement a triplicate of an BSA standard curve was used to convert intensities to concentration. For each measurement a triplicate of a total concentration (c_tot_) series below and above the saturation concertation (c_dil_) were carried out. Below c_dil_ we see a match of c_tot_ and the measured concentrations (c_BCA_) and as expected above c_dil_ a plateau-like region for c_BCA_. The intersect of linear fits to the plateau and the initial slope provides us with c_dil_ (Extended Data Fig. 8).

To determine the condensed phase concentration c_con_ via fluorescence absorption measurement with a NanoDrop spectrometer (ThermoFisher, USA) we used a negative displacement pipette (Eppendorf, Germany). This was necessary because of the high viscosity of the condensed phase and potential errors using air pipettes. A known volume of condensed phase was diluted into high salt buffer before absorption measurement to ensure dissolution of the phase. In addition, guanidinium chloride can be used to dissolve condensates for proteins that would not dissolve again in high salt buffer.

### Temperature shift experiments

Emulsions were prepared as described in the section “Preparation of emulsions” except without a final centrifugation step. This step was not necessary since the total protein concentrations were adjusted to be outside the binodal at the temperatures used for preparation. We used FUS::GFP at the salt concentrations of 100, 150, 200, and 300 mM KCl at protein concentrations of 2.4, 4.8, 7.9, and 15.9 µM respectively (all pH 7.4). A low density of emulsion droplets was mounted in Parafilm chambers on the temperature-controlled stage to ensure stationary emulsions during temperature shift experiments. Images were recorded using the described spinning disc confocal system at 10 s temporal resolution and 1 µm resolution in the z-axis using a 40x air objective.

### RNA experiments

Experiments were prepared as described above for binary systems however the ratio between RNA and protein concentration was kept constant for the titrations of total protein concentration. We used poly(A) RNA (P9403, Sigma-Aldrich), buffers were prepared using nuclease free water (AM9938, Invitrogen), and surfaces were cleaned with RNaseZap (AM9780, Invitrogen). The RNA concentration was calculated using the average molecular weight reported in the product documentation. While the dilute protein concentration c_dil_ should be accurately represented, the condensed concentration c_con_ represents and effective value that is made up of both RNA and protein concentration in the condensed phase.

### One-cell *C. elegans* embryo experiments

One-cell embryos were imaged and analysed according to a protocol described previously^16^. This allowed us to derive the volume fraction of condensed phase as well as the total concentration of labelled protein per embryo. In brief, known total protein concentrations (PGL-1::GFP, PGL-3::GFP respectively) in the different worm strains were related to fluorescence intensities to derive a scaling factor between florescence intensity and concentration. To derive the total condensate volume as well as the embryo volume we used a custom-written MATLAB (The MathWorks) segmentation routine. The worm lines used were TH586, pgl-1::mEGFP pgl-1(dd54[pgl-1::mEGFP]) and TH561, pgl-3::mEGFP (pgl-3(dd29[pgl-3::mEGFP]). Here, the proteins PGL-1 and PGL-3 were labelled at their endogenous genomic locus with monomeric enhanced GFP using the co-CRISPR method.

### PGL-3 protein purification

PGL-3 was purified from insect cells according to Saha et al.^5^ from SF9-ESF cells were infected with baculovirus containing the PGL-3-GFP-6HIS protein under the polyhedrin promoter. Cells were harvested after 3 days of infection by centrifugation at 500 x g for 10 min and then resuspended in lysis buffer (25 mM HEPES 7.25, 300 mM KCl, 10 mM imidazole, 1 mM DTT, 1 protease inhibitor). Cells were lysed by passing the cells 2 times through the LM20 microfluidizer at 15000 psi. The lysate was then centrifuged at 20000 rpm for 45 min at 15 °C. The lysate was loaded in a pre-equilibrated Ni-NTA column with lysis buffer at 3 mL/min. The Ni-NTA column was rinsed with 10 C.V of wash buffer (25 mM HEPES 7.25, 300 mM KCl, 20 mM imidazole, 1 mM DTT, 1) and the protein was eluted in 1.5 mL fractions with elution buffer (25 mM HEPES 7.25, 300 mM KCl, 250 mM imidazole, 1 mM DTT). After elution the GFP tagged was cleaved to produce untagged PGL-3. The cleavage was performed using a TEV protease overnight at 4 °C. PGL-3 and PGL-3-GFP proteins were diluted with Dilution buffer (25 mM Tris pH 8.0, 1 mM DTT) to reach 50 mM KCl before loading the protein in an anion exchange HiTrapQ HP 5 mL column. The HiTrap column was previously equilibrated first with HiTrapQ elution buffer (25 mM Tris pH 8.0, 50 mM KCl, 1 mM DTT) and then with HiTrapQ binding buffer (25 mM Tris pH 8.0, 1 M KCl, 1 mM DTT). The column was mounted in a Äkta Pure FPLC system. After the sample was loaded the column was washed with HiTrapQ binding buffer. The sample was finally eluted with a linear gradient from 0 to 55% of HiTrapQ elution buffer (25 mM Tris pH 8.0, 1 M KCl 1 mM DTT) for 25 C.V. Finally, a 100% HiTrap elution buffer step was performed for 5 C.V. The pooled fractions were then loaded in a HiLoad 16/60 Superdex 200 size exclusion chromatography column that was previously equilibrated with superdex buffer (25 mM HEPES 7.25, 300 mM KCl, 1 mM DTT). After size exclusion, the final samples were collected.

### FUS protein purification

FUS-GFP was purified as previously described in Wang et al. ^10^.

## Acknowledgements

For fruitful discussions and help with protein work we thank, Martine Ruer-Gruß, Patrick McCall, Tylor Harmon, Jie Wang, Christoph Weber, Esteban Meca, Barbara Wagner, Simone Reber, and Titus Franzmann. For comments on the manuscript, we would like to thank Sina Wittmann, Carsten Hoege, Cristina Jiménez-López, and André Nadler. We thank the light microscopy, chromatography, and protein purification facilities at MPI-CBG for their support with this project. We thank Olympus for providing the CSU-W1 SoRa spinning-disc system based on an IXplore IX83 microscope. A.W.F. was supported by the ELBE postdoctoral fellows program and the Max Planck Research Network for Synthetic Biology (MaxSynBio) consortium, jointly funded by the Federal Ministry of Education and Research of Germany and the Max Planck Society. A.A.H acknowledges support from the NOMIS foundation.

## Author contributions

A.W.F, J.M.I.A and A.A.H. planned the research and wrote the manuscript. A.W.F, J.M.I.A conceived and performed the research, wrote the code used in the analysis and evaluated the data.

## Extended Data

**Extended Data Figure 1:**
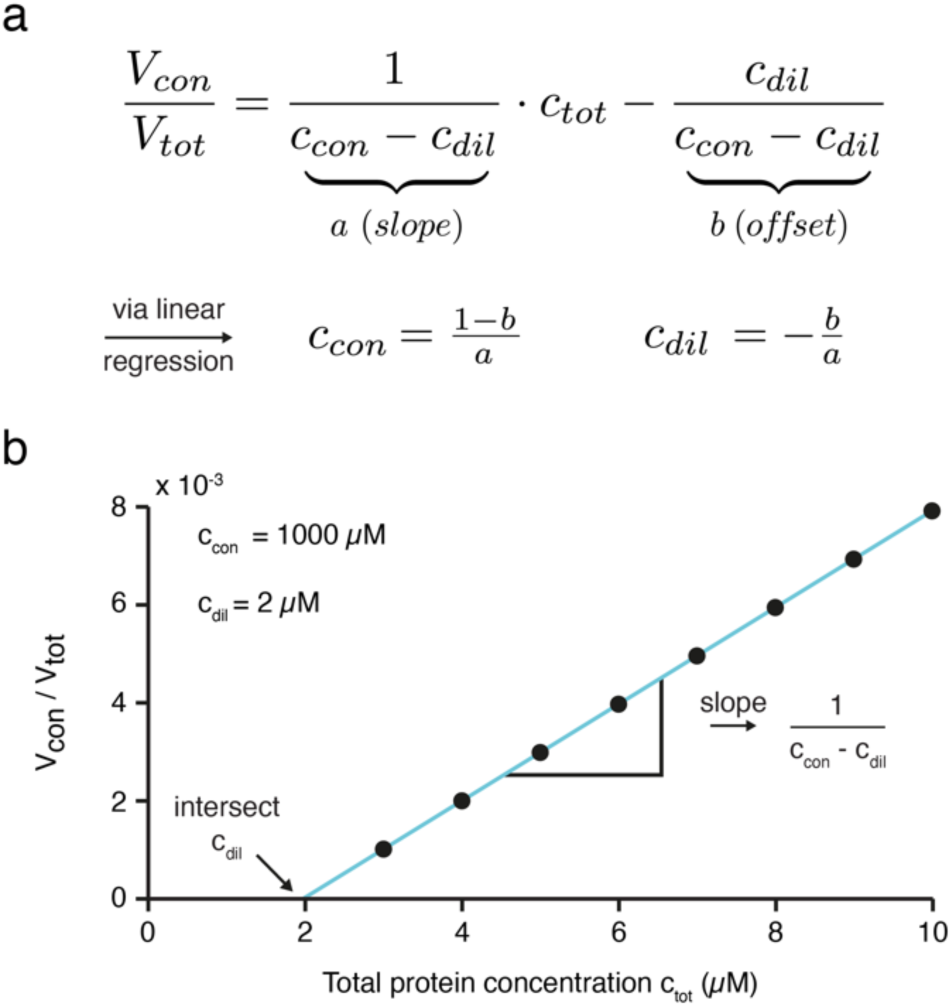
Principle of inPhase. **a,** Central equation derived from volume and mass conversation. A linear regression of the volume fraction of condensed phase against total protein concentration is used to estimate c_con_ and c_dil_ via the slope a and offset b. **b**, Example graph of volume fraction versus total protein concentration for a fictitious protein with c_dil_ = 2 µM and c_con_ = 1000 µM.

**Extended Data Figure 2:**
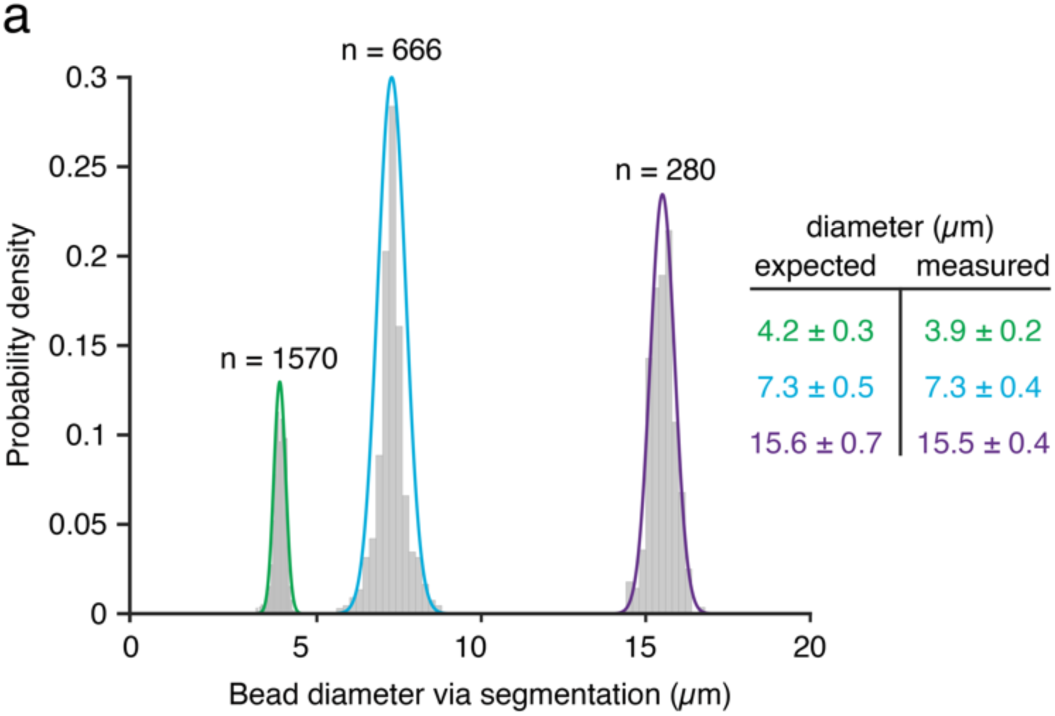
Segmentation correctly estimates the diameter of standard beads. **a**, Diameter of fluorescent polystyrene beads as determined by our segmentation script in comparison to the specifications given by the manufacturer.

**Extended Data Figure 3:**
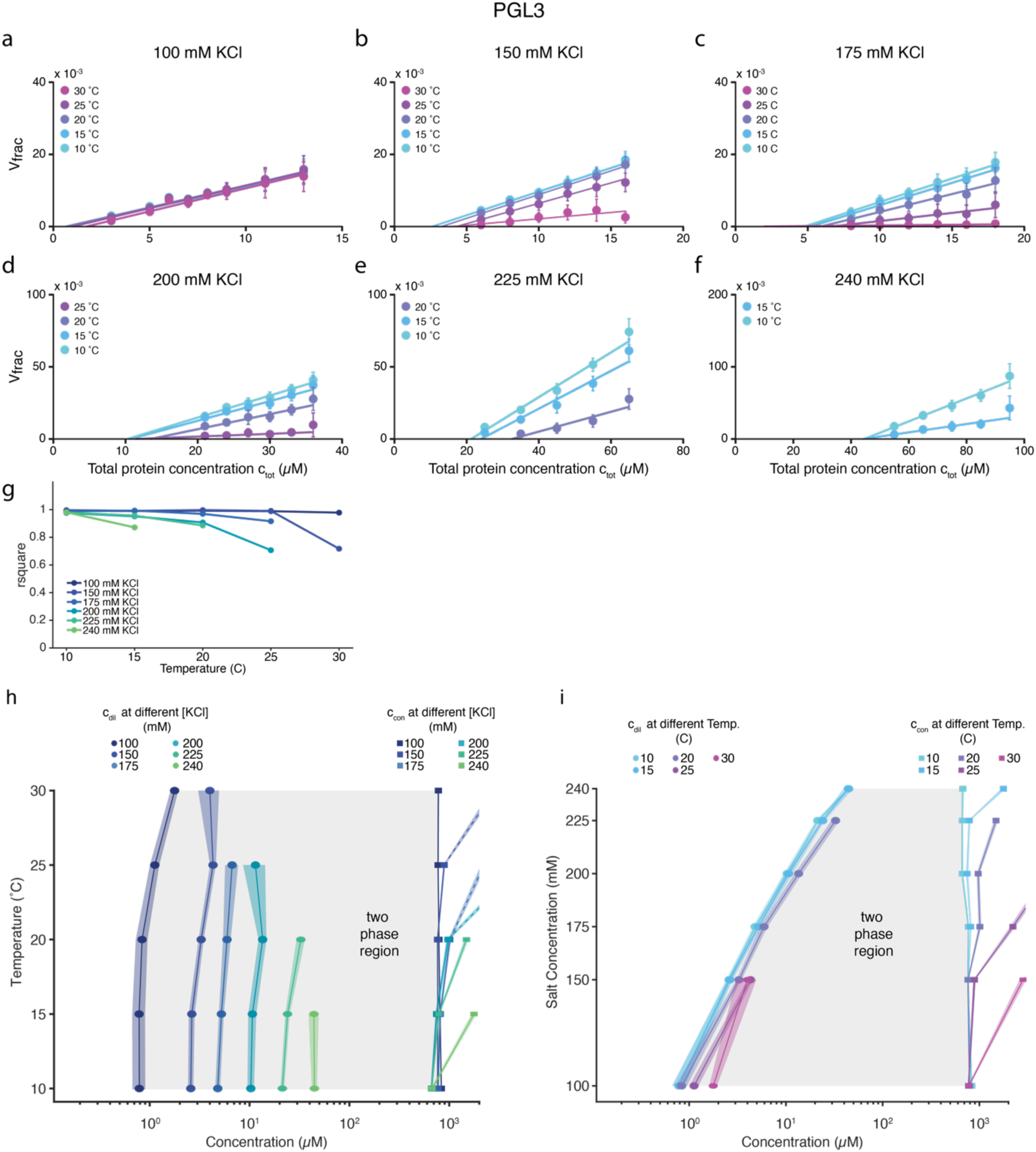
Linear regressions of V_frac_(c_tot_) data and phase diagrams for PGL-3. **a-f**, Linear regressions of V_frac_(c_tot_) of samples prepared at 100 (a), 150 (b), 175 (c), 200 (d) and 225 (e) mM KCl and measured at different temperatures. Experimental data is pooled from all independent repeats and then fitted to Eq. 1. Each individual experiment consists of datapoints from many emulsion droplets. Error bars depict standard deviations. **g**, Quality of linear regression (R-squared value) of the data presented in panel a-f. The R-squared value was obtained by fitting Eq. 1 to the mean V_frac_ at each protein concentration. **h, i**, Temperature (h) and salt (i) phase diagrams presenting the derived data for c_con_ and c_dil_ from the linear regressions in panel a-f. Lines connecting the data points are a guide to the eye. Shaded areas depict the SD derived by error propagation of the standard deviation of the linear regressions in panel a-f. See Supplementary Table 1 for the number of repeats.

**Extended Data Figure 4:**
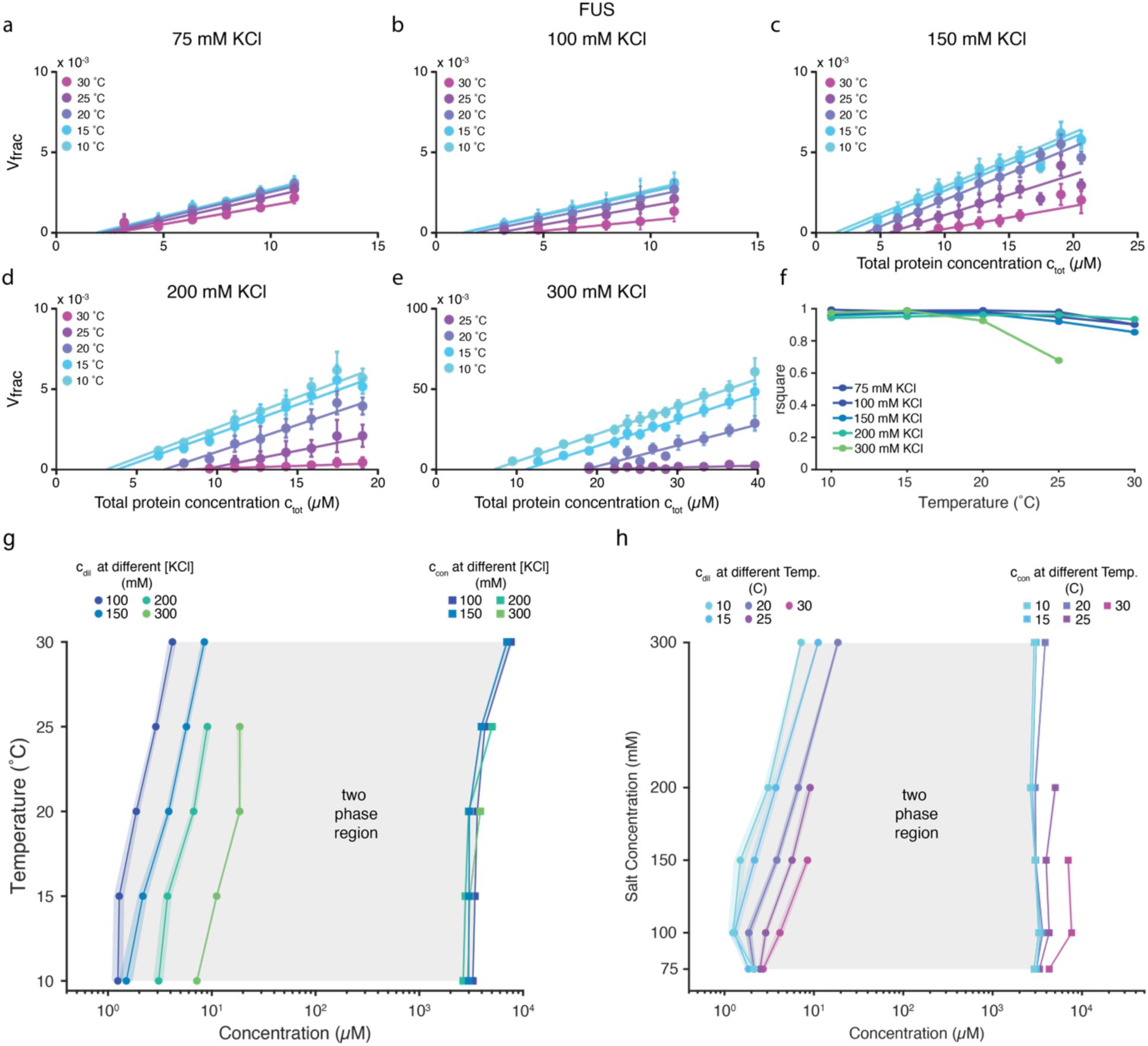
Linear regressions of V_frac_(c_tot_) data and phase diagrams for FUS. **a-e**, Linear regressions of V_frac_(c_tot_) of samples prepared at 75 (a), 100 (b), 150 (c), 200 (d) and 300 (e) mM KCl and measured at different temperatures. Experimental data is pooled from all independent repeats and then fitted to Eq. 1. Each individual experiment consists of data points from many emulsion droplets. Error bars depict standard deviations. **f**, Quality of linear regression (R-squared value) of data presented in panel a-e. The R-squared value was obtained by fitting Eq. 1 to the mean V_frac_ at each protein concentration. **g**, **h**, Temperature (g) and salt (h) phase diagrams presenting the derived data for c_con_ and c_dil_ from the linear regressions in panel a-e. Lines connecting the data points are a guide to the eye. Shaded areas depict the errors derived by error propagation of the SD of the linear regression in panel a-e. See Supplementary Table 2 for the number of repeats.

**Extended Data Figure 5:**
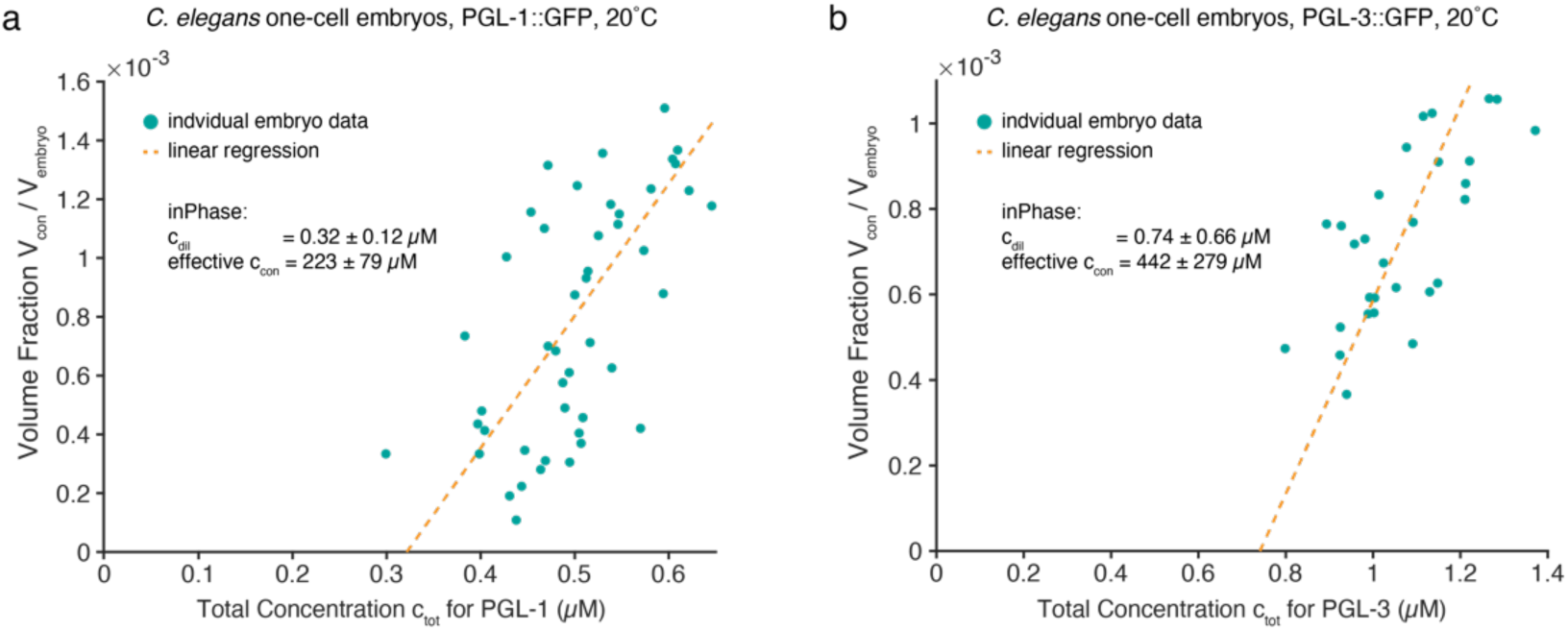
Protein concentration heterogeneity in C. elegans embryos allows to test inPhase in vivo. **a**, Data of condensed volume fraction against total PGL-1::GFP concentration at 20°C (green dots, N=46). Linear regression to the data (dashed orange line) allows to use equation 1 and derive c_dil_ and effective c_cond_ concentrations. **b**, Data of condensed volume fraction against total PGL-3::GFP concentration at 20°C (green dots, N=29). Linear regression to the data (dashed orange line) allows to use equation 1 and derive c_dil_ and effective c_cond_ concentrations.

**Extended Data Figure 6:**
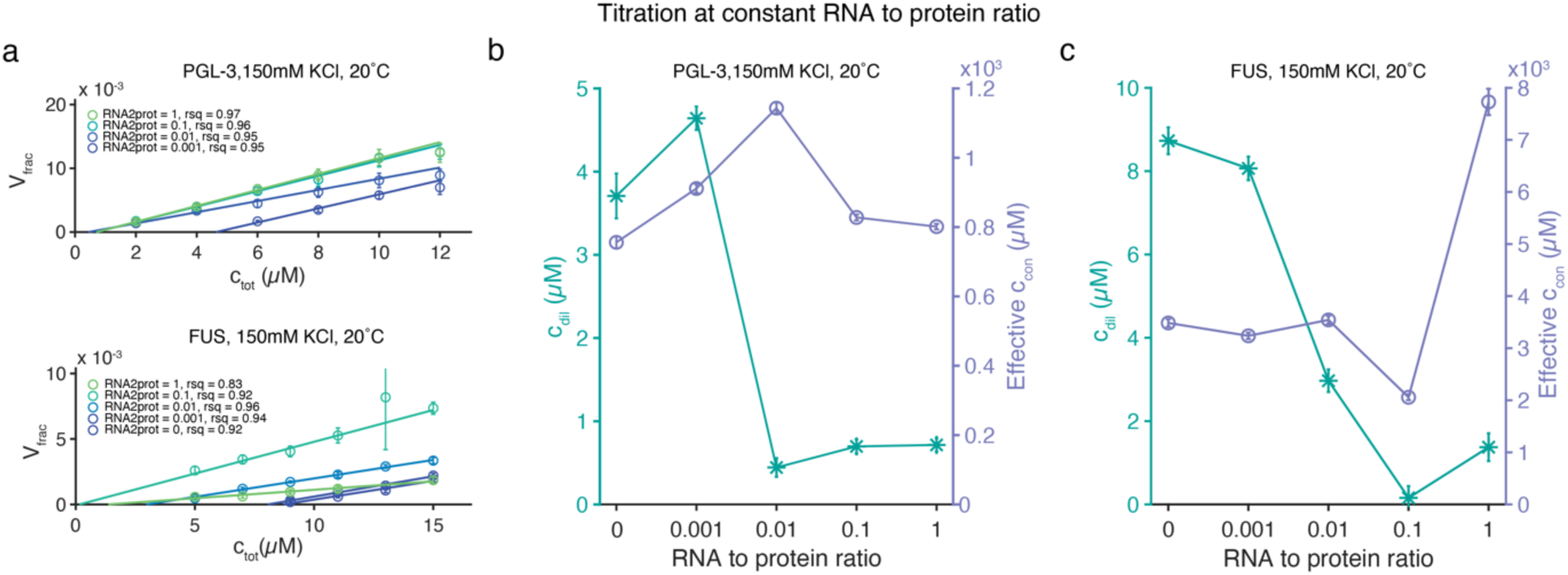
Increasing the RNA to protein ratio reduces c_dil_. **a**, Titration of total protein concentration at a constant ratio of poly(A) RNA to protein concentration. R-squared values are from linear regression to the individual curves (errors depict standard deviation). **b-c**, Derived values of c_dil_ and effective c_con_ using inPhase on the data presented in panel a.

**Extended Data Figure 7:**
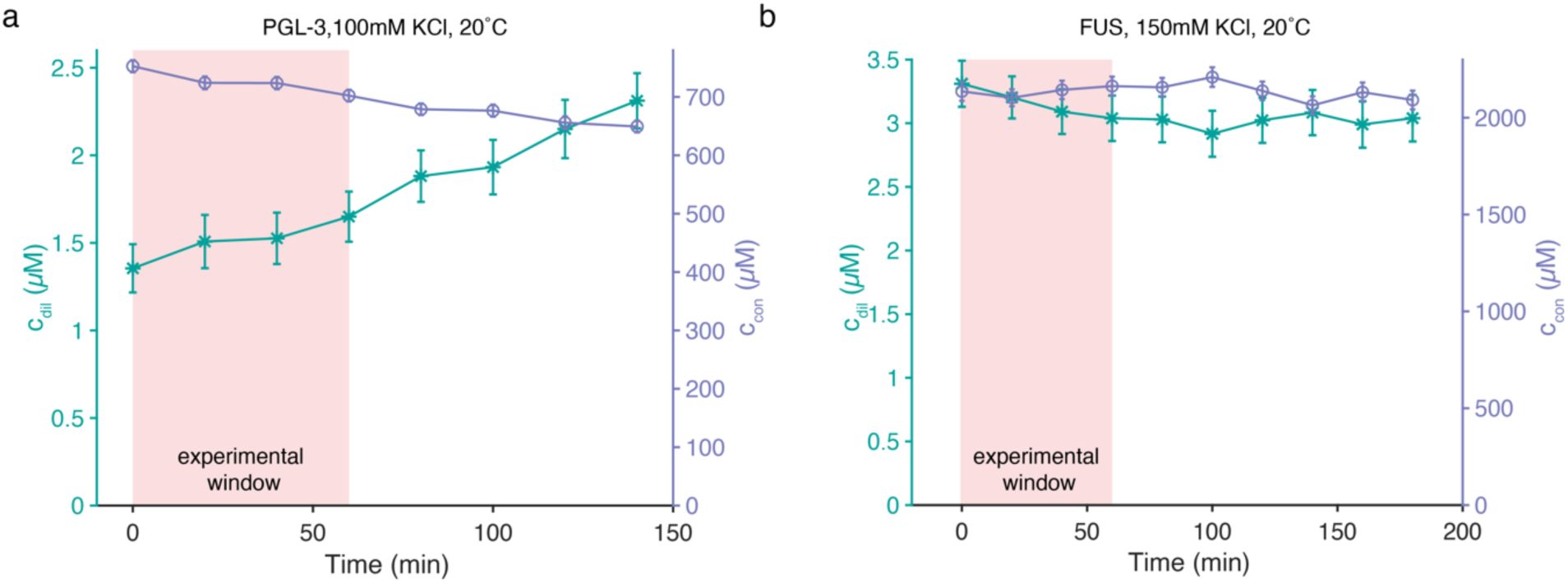
c_dil_ and c_con_ do not change considerably during the experimental window. **a**, **b**, Time dependent inPhase evaluation of dilute and condensed branch concentrations for PGL-3 (a) and FUS (b) (Errors from error propagation of linear regression confidence intervals). Shaded areas depict the time-window used for the temperature and salt dependent phase diagrams.

**Extended Data Figure 8:**
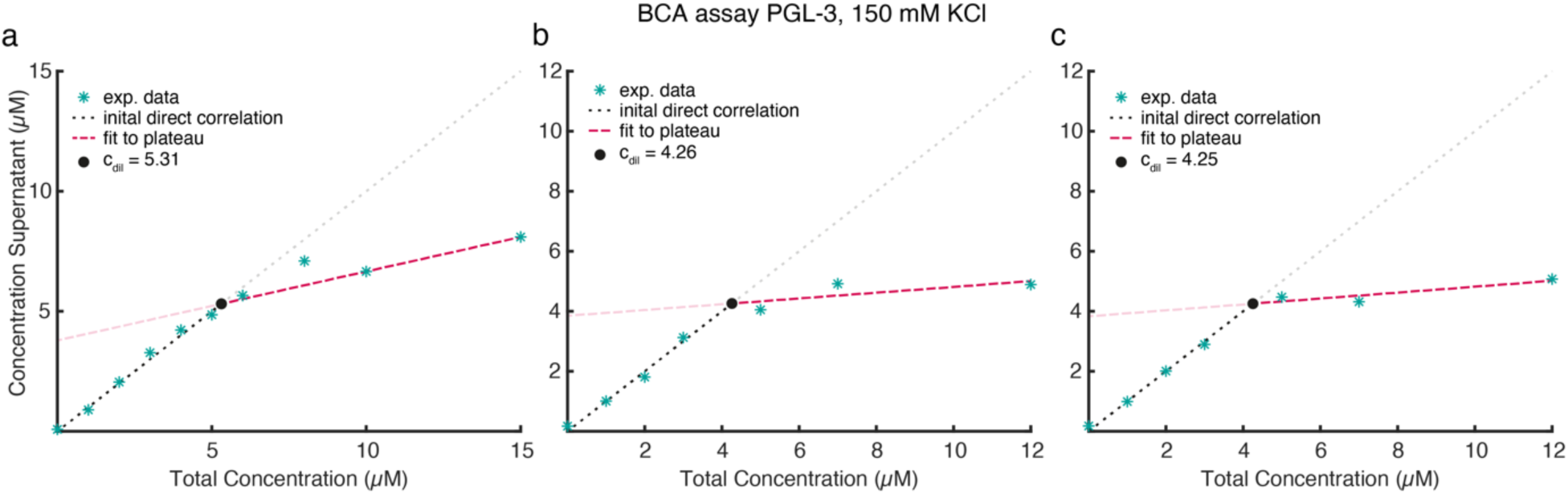
Bicinchoninic acid (BCA) assay to determine c_dil_. **a-c**, Three repeats of the measured supernatant concentrations versus total protein concentration used for the assay. Intersect of initial direct correlation between input total and measured supernatant concentrations and the plateau-like region yields c_dil_. Datapoints are mean values of technical triplicates

## Notes

https://git.mpi-cbg.de/fritsch/inphase-code-repository

### Competing Interest Statement

A.A.H. is cofounder and member of the scientific advisory board of Dewpoint Therapeutics. A.W.F., J.M.I.A., and A.A.H. have filed a patent application on the presented method.

### Summary of Updates

Acknowledgement section updated to include the NOMIS foundation

