## Supplementary Information for "inPhase — A simple, accurate and fast approach to determine phase diagrams of protein condensates"

**Supplementary Video 1**:
Maximum projection of one emulsion droplet containing FUS:GFP at a concentration of 2.4 µM at 100 mM KCl and a pH of 7.4. Time is indicated in the top left corner in mm:ss. The temperature is indicated in the top right corner. Scale bar depicts 5 µM.

**Supplementary Video 2**:
Maximum projection of one emulsion droplet containing FUS:GFP at a concentration of 4.8 µM at 150 mM KCl and a pH of 7.4. Time is indicated in the top left corner in mm:ss. The temperature is indicated in the top right corner. Scale bar depicts 5 µM.

**Supplementary Video 3**:
Maximum projection of one emulsion droplet containing FUS:GFP at a concentration of 7.9 µM at 200 mM KCl and a pH of 7.4. Time is indicated in the top left corner in mm:ss. The temperature is indicated in the top right corner. Scale bar depicts 5 µM.

**Supplementary Video 4**:
Maximum projection of one emulsion droplet containing FUS:GFP at a concentration of 15.9 µM at 300 mM KCl and a pH of 7.4. Time is indicated in the top left corner in mm:ss. The temperature is indicated in the top right corner. Scale bar depicts 5 µM.

**Supplementary Table 1:**
Data used for PGL-3 phase diagrams and partition factor comparison.

| c Salt (mM KCl) | T (˚C) | Label Frac. | pH | n emulsion total | average n emulsion per $c_{tot}$ | distinct repeat | $c_{dil}$ (µM) | $c_{dil}$ SD (µM) | $c_{con}$ (µM) | $c_{con}$ SD (µM) | PF inPhase | PF intensity | $R^2$ |
|---|---|---|---|---|---|---|---|---|---|---|---|---|---|
| **100** | 10 | 0.05 | 7.4 | 626 | 78 | 3 | 0.79 | 0.12 | 834 | 12 | 1062 | 157 | 0.992 |
| **100** | 15 | 0.05 | 7.4 | 648 | 81 | 3 | 0.78 | 0.11 | 808 | 12 | 1034 | 193 | 0.990 |
| **100** | 20 | 0.05 | 7.4 | 627 | 78 | 3 | 0.84 | 0.11 | 789 | 11 | 943 | 201 | 0.992 |
| **100** | 25 | 0.05 | 7.4 | 621 | 77 | 3 | 1.12 | 0.12 | 773 | 12 | 688 | 182 | 0.989 |
| **100** | 30 | 0.05 | 7.4 | 636 | 79 | 3 | 1.77 | 0.16 | 778 | 15 | 439 | 138 | 0.978 |
| **150** | 10 | 0.05 | 7.4 | 552 | 92 | 3 | 2.57 | 0.16 | 763 | 10 | 297 | 169 | 0.996 |
| **150** | 15 | 0.05 | 7.4 | 520 | 86 | 3 | 2.64 | 0.18 | 769 | 12 | 292 | 160 | 0.991 |
| **150** | 20 | 0.05 | 7.4 | 519 | 86 | 3 | 3.27 | 0.19 | 764 | 13 | 233 | 128 | 0.997 |
| **150** | 25 | 0.05 | 7.4 | 537 | 89 | 3 | 4.30 | 0.29 | 890 | 22 | 207 | 91 | 0.991 |
| **150** | 30 | 0.05 | 7.4 | 480 | 80 | 3 | 4.00 | 0.94 | 2826 | 225 | 707 | 60 | 0.717 |
| **175** | 10 | 0.05 | 7.4 | 553 | 92 | 3 | 4.80 | 0.27 | 772 | 14 | 161 | 109 | 0.988 |
| **175** | 15 | 0.05 | 7.4 | 570 | 95 | 3 | 5.22 | 0.29 | 813 | 16 | 156 | 94 | 0.993 |
| **175** | 20 | 0.05 | 7.4 | 555 | 92 | 3 | 5.94 | 0.51 | 1013 | 35 | 170 | 72 | 0.969 |
| **175** | 25 | 0.05 | 7.4 | 534 | 89 | 3 | 6.69 | 0.99 | 2222 | 145 | 332 | 53 | 0.916 |
| **200** | 10 | 0.05 | 7.4 | 553 | 92 | 3 | 10.29 | 0.77 | 666 | 16 | 65 | 49 | 0.976 |
| **200** | 15 | 0.05 | 7.4 | 541 | 90 | 3 | 10.68 | 0.98 | 748 | 23 | 70 | 41 | 0.952 |
| **200** | 20 | 0.05 | 7.4 | 537 | 89 | 3 | 13.53 | 1.22 | 972 | 35 | 72 | 29 | 0.907 |
| **200** | 25 | 0.05 | 7.4 | 515 | 85 | 3 | 11.46 | 2.84 | 5235 | 453 | 457 | 22 | 0.707 |
| **225** | 10 | 0.05 | 7.4 | 196 | 39 | 1 | 21.29 | 0.92 | 666 | 12 | 31 | 28 | 0.980 |
| **225** | 15 | 0.05 | 7.4 | 197 | 39 | 1 | 23.99 | 1.21 | 787 | 18 | 33 | 22 | 0.957 |
| **225** | 20 | 0.05 | 7.4 | 161 | 40 | 1 | 32.61 | 3.06 | 1487 | 75 | 46 | 17 | 0.886 |
| **240** | 10 | 0.05 | 7.4 | 159 | 31 | 1 | 44.61 | 2.54 | 675 | 20 | 15 | 10 | 0.981 |
| **240** | 15 | 0.05 | 7.4 | 161 | 32 | 1 | 44.10 | 5.02 | 1774 | 103 | 40 | 8 | 0.872 |

**Label Frac.**: Amount of GFP fused protein relative to the untagged version.
**n emulsion total**: Sum of all emulsion droplets pooled from all repeats.
**average n emulsion per $c_{tot}$**: Average number of emulsion droplets used per total concentration in the titration series over all repeats.
**$c_{dil}$**: Derived from the fit of equation 1 (main manuscript) to the combined volume fraction data, using all individual emulsion droplets from all repeats. Fit was performed using robust bi-square routine.
**$c_{dil}$ SD**: Derived via error propagation of the SD of the fit of equation 1 (main manuscript) to the combined volume fraction data, using all individual emulsion droplets from all repeats.
**$c_{con}$**: See $c_{dil}$.
**$c_{con}$ SD**: See $c_{dil}$ SD.
**PF inPhase**: Partition factor derived by division of $c_{con}$ / $c_{dil}$.
**PF intensity**: Mean partition factor derived by division of fluorescence intensity values $I_{con}$ / $I_{dil}$ using only condensates above a threshold size.
**$R^2$**: R square value of the fit of equation 1 (main manuscript) to the mean volume fraction values derived from the pooled data of all repeats weighted by emulsion droplet numbers.

**Supplementary Table 2:**

Data used for FUS phase diagrams and partition factor comparison.

| c Salt (mM KCl) | T (˚C) | Label Frac. | pH | n emulsion total | average n emulsion per $c_{tot}$ | distinct repeat | $c_{dil}$ (µM) | $c_{dil}$ SD (µM) | $c_{con}$ (µM) | $c_{con}$ SD (µM) | PF inPhase | PF intensity | $R^2$ |
|---|---|---|---|---|---|---|---|---|---|---|---|---|---|
| **75** | 10 | 1 | 7.4 | 423 | 70 | 2 | 2.14 | 0.15 | 2916 | 53 | 1361 | 550 | 0.994 |
| **75** | 15 | 1 | 7.4 | 416 | 69 | 2 | 1.85 | 0.15 | 3085 | 57 | 1665 | 512 | 0.982 |
| **75** | 20 | 1 | 7.4 | 417 | 69 | 2 | 2.14 | 0.13 | 3096 | 51 | 1446 | 438 | 0.978 |
| **75** | 25 | 1 | 7.4 | 390 | 65 | 2 | 2.50 | 0.14 | 3343 | 56 | 1339 | 338 | 0.949 |
| **75** | 30 | 1 | 7.4 | 394 | 65 | 2 | 2.70 | 0.17 | 4297 | 86 | 1590 | 248 | 0.901 |
| **100** | 10 | 1 | 7.4 | 725 | 120 | 5 | 1.24 | 0.16 | 3285 | 69 | 2642 | 682 | 0.960 |
| **100** | 15 | 1 | 7.4 | 719 | 119 | 5 | 1.28 | 0.15 | 3438 | 68 | 2681 | 632 | 0.989 |
| **100** | 20 | 1 | 7.4 | 690 | 115 | 5 | 1.87 | 0.16 | 3624 | 74 | 1936 | 508 | 0.990 |
| **100** | 25 | 1 | 7.4 | 710 | 118 | 5 | 2.89 | 0.18 | 4289 | 96 | 1486 | 354 | 0.980 |
| **100** | 30 | 1 | 7.4 | 503 | 100 | 4 | 4.17 | 0.41 | 7654 | 344 | 1834 | 175 | 0.902 |
| **150** | 10 | 1 | 7.4 | 653 | 59 | 4 | 1.50 | 0.23 | 2985 | 54 | 1990 | 565 | 0.958 |
| **150** | 15 | 1 | 7.4 | 660 | 60 | 4 | 2.17 | 0.24 | 3016 | 56 | 1392 | 442 | 0.972 |
| **150** | 20 | 1 | 7.4 | 662 | 60 | 4 | 3.85 | 0.26 | 3038 | 60 | 790 | 303 | 0.970 |
| **150** | 25 | 1 | 7.4 | 641 | 64 | 4 | 5.70 | 0.37 | 3973 | 104 | 697 | nan | 0.920 |
| **150** | 30 | 1 | 7.4 | 526 | 75 | 4 | 8.47 | 0.59 | 7009 | 264 | 827 | nan | 0.854 |
| **200** | 10 | 1 | 7.4 | 397 | 44 | 3 | 3.07 | 0.36 | 2650 | 63 | 862 | 427 | 0.943 |
| **200** | 15 | 1 | 7.4 | 410 | 45 | 3 | 3.74 | 0.38 | 2785 | 70 | 744 | 333 | 0.952 |
| **200** | 20 | 1 | 7.4 | 394 | 49 | 3 | 6.68 | 0.46 | 2990 | 85 | 448 | 230 | 0.961 |
| **200** | 25 | 1 | 7.4 | 399 | 57 | 3 | 9.07 | 0.80 | 5025 | 229 | 554 | 124 | 0.962 |
| **300** | 10 | 1 | 7.4 | 1038 | 79 | 5 | 7.19 | 0.19 | 2925 | 21 | 407 | 249 | 0.974 |
| **300** | 15 | 1 | 7.4 | 1023 | 85 | 5 | 11.15 | 0.24 | 3070 | 26 | 275 | 168 | 0.987 |
| **300** | 20 | 1 | 7.4 | 818 | 81 | 5 | 18.55 | 0.56 | 3868 | 64 | 209 | 88 | 0.925 |

For a description of the columns see Supplementary Table 1.

**Supplementary Table 3:**

Data used to study the influence of the fluorescence label fraction phase separation and quantitative fluorescence analysis.

| Label Frac. | c Salt (mM KCl) | T (°C) | pH | n emulsion total | average n emulsion per $c_{tot}$ | distinct repeat | $c_{dil}$ (µM) | $c_{dil}$ SD (µM) | $c_{con}$ (µM) | $c_{con}$ SD (µM) | PF inPhase | PF intensity | $R^2$ |
|---|---|---|---|---|---|---|---|---|---|---|---|---|---|
| 0.002 | 100 | 20 | 7 | 609 | 55 | 4 | 1.26 | 0.10 | 834 | 10 | 661 | 147 | 0.972 |
| 0.01 | 100 | 20 | 7 | 983 | 89 | 5 | 1.28 | 0.07 | 804 | 7 | 630 | 187 | 0.973 |
| 0.05 | 100 | 20 | 7 | 574 | 95 | 3 | 0.89 | 0.06 | 870 | 7 | 973 | 245 | 0.996 |
| 0.2 | 100 | 20 | 7 | 530 | 53 | 3 | 1.10 | 0.08 | 809 | 9 | 733 | 212 | 0.977 |
| 0.5 | 100 | 20 | 7 | 578 | 52 | 3 | 0.75 | 0.13 | 845 | 14 | 1133 | 247 | 0.970 |
| 0.75 | 100 | 20 | 7 | 667 | 95 | 4 | 0.62 | 0.08 | 923 | 10 | 1494 | 246 | 0.986 |
| 1 | 100 | 20 | 7 | 765 | 127 | 4 | 0.72 | 0.14 | 803 | 14 | 1113 | 225 | 0.994 |

For a description of the columns see Supplementary Table 1.
